# Bioglass-embedded alginate scaffold for dental bone tissue engineering applications

**DOI:** 10.64898/2026.01.12.698929

**Authors:** Ananya N Nayak, Roshni Ramachandran

**Affiliations:** Department of Biotechnology, Ramaiah Institute of Technology, Bengaluru, Karnataka -560054, India

## Abstract

Traumatic dental injury impacts approximately 1 in 10 people globally and it is further challenging to treat with the complications such as inflammation, infection and dynamic oral pH which collectively slow down the dental bone healing. Currently existing approaches address these complications individually which results in suboptimal regenerative outcomes. Our study focuses on developing a multifunctional dental scaffold engineered to tackle inflammation and infection simultaneously and enhance bone regeneration through a dual drug delivery strategy and integration of Bioglass.

A novel citric acid mediated process was used to produce bioglass which was further characterized using XRD and SEM analyses. The bioglass was capped with antibiotic and integrated into the alginate scaffold which was further subjected to surface coating of painkiller to enable rapid anti-inflammatory action and sustained antimicrobial release. The composite scaffold was further assessed for its physiochemical properties using swelling and degradation analysis, SEM was carried out to understand the structure and morphology of the scaffold. MTT assays were carried out on osteoblastic and fibroblastic cell lines to understand the cytocompatibility of the scaffold, while the osteogenic property was evaluated through biomineralization assay.

The results showed successful synthesis and homogenous integration of bioglass, leading to increased swelling potential, controlled degradation and excellent biocompatibility. Robust osteogenic differentiation validated the scaffold’s capacity as an advanced platform for dental bone tissue engineering and effective management of traumatic dental injuries.

**Graphical abstract:** 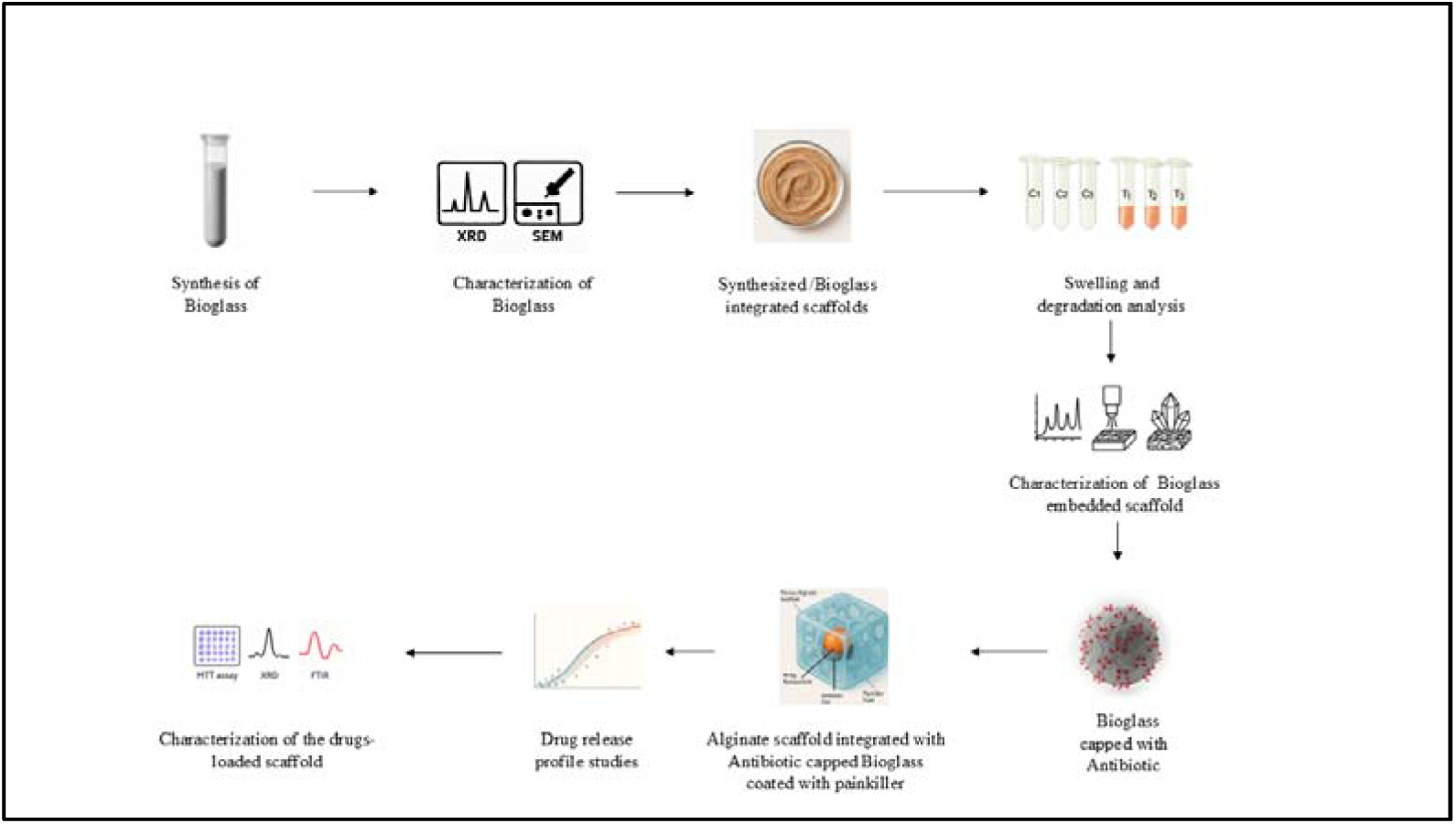

## 1) Introduction

The emergence of nanocomposites has opened new possibilities in dental bone tissue engineering. Studies have demonstrated the promise of alginate/hydroxyapatite scaffolds in promoting osteogenic differentiation of dental pulp stem cells, which has implications for alveolar bone regeneration (**Sancilio et al., 2018**). One of the works on 3D-printed PLGA/TiO2 nanocomposites showed enhanced osteoblast activity, indicating improved scaffold designs for bone regeneration(**Rasoulianboroujeniet al., 2019**). These studies, along with others, highlight the potential of nanocomposites in improving mechanical properties, biocompatibility, and bone regeneration capabilities in the dental context(**Ferrer et al 2020, Albalwi et al. 2022**).

Past researchers have explored bioglass for tissue engineering and bone regeneration. Bioglass possesses unique properties that make it a promising material for bone regeneration, including the ability to form a hydroxyapatite layer for integration with bone tissue, support bone growth through osteoconductivity, and release ions that enhance cellular activity. Additionally, its customizable composition allows for tailoring mechanical properties and bioactivity to specific applications (**Ribas et al., 2019**). Studies have explored various scaffold types, including three-dimensional membranous scaffolds composed of bioglass and alginate composites(**Bargavi et al., 2020**). In addition, bioglass-alginate composite hydrogel beads have been investigated as cell carriers for bone regeneration(**Zeng et al., 2014**). Another study focused on alginate-bioactive glass containing Zn and Mg composite scaffolds for bone tissue engineering, examining their mechanical properties, in vitro behavior, cytotoxicity, cell behavior, ion release, and antibacterial activity(**Zamani et al., 2019)**. Together, these studies highlight the diverse applications of bioglass in bone regeneration and the importance of in vitro and in vivo evaluations to determine their efficacy and safety.

### 2) Methodology

#### 2.1) Synthesis and Characterization of Bioglass

An aqueous solution of citric acid was prepared. Silica gel and ammonium dihydrogen orthophosphate were added to the mixture with constant stirring unless a clear solution was obtained. Sodium nitrate and Calcium nitrate tetrahydrate were added to the mixture at constant stirring unless a homogenous mixture was obtained. The mixture was left to age for 24 hours, after which the mixture attained a gel-like consistency. The gel was subjected to drying for 12h at 60□C until the gel dried up completely. Further, it was subjected to drying at 200□C for 5 hours, followed by calcination at 700□C for 2 hours. For qualitative analysis, XRD patterns were recorded in the interval 10□ < 2θ >800**(Faure et al., 2015)**. SEM analysis was carried out to understand the morphology of the nanoparticle. The Bioglass was subjected to gold sputtering for two minutes using DII-29030SCTRSmart coater, followed by SEM-EDS analysis using Ultra55 FE-SEM Carl Zeiss EDS.

#### 2.2) Preparation and characterization of Bioglass embedded scaffold

Sodium alginate (1. 6 g) was dispersed in 20 mL of water and stirred magnetically until a uniform solution was achieved. Subsequently, 10 mg of the synthesized bioglass was added to the alginate solution and stirred continuously to ensure a homogeneous mixture. The alginate-bioglass mixture was then added dropwise to a chilled calcium chloride solution to induce gel formation. The resulting gel was immediately frozen at -20°C and subsequently freeze-dried to yield a porous scaffold. The scaffold was characterized using SEM analysis to understand the scaffold’s morphology, XRD, and FTIR to understand if the chemical integrity of alginate and the scaffold is maintained in the scaffold.

#### 2.3) Swelling and degradation analysis of bioglass-embedded scaffolds

##### 2.3.1) Swelling assay for bioglass-embedded scaffold

The nanocomposite scaffold was added into a known volume of PBS for 21 days The Wet weight of the scaffolds were measured at , 1, 3, 5, 7, 9, 11, 13, 15, 17, 19, and 21 days respectively to calculate the swelling ratio as follows:

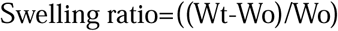

Wherein, Wo is the initial dry weight of the scaffold, Wt is the wet weight of the scaffold on day T.

##### 2.3.2) Degradation assay for bioglass-embedded scaffold

The Dry weight of the nanocomposite scaffold was noted. The scaffold was added to the known volume of PBS and trypsin mixture for 15 days . Wet weight of the scaffolds were measured on days 1, 3, 5, 7, 9, 11, 15 days respectively degradation ratio was calculated as follows:

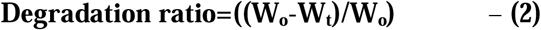

Wherein W_t_ is the wet weight of the swollen scaffold, W_o_ is the initial weight of the dry scaffold.

#### 2.4) Biomineralization studies

A 100 mL solution of simulated body fluid (SBF) was prepared using the following reagents: 0. 745 g NaCl, 0. 044 g KCl, 0. 035 g NaHCO3, 0. 116 g CaCl2·2H2O, 0. 031 g MgSO4·7H2O, 0. 023 g Na2HPO4·2H2O, and 0. 15 g Tris base. These reagents were dissolved in 90 mL of deionized water under constant magnetic stirring. The pH of the solution was adjusted to 7. 4 using 1 M HCl. Subsequently, the solution was brought to a final volume of 100 mL with deionized water(**Bellucci et al., 2019**).

##### Treatment of Scaffolds and Analysis

Scaffolds embedded with bioglass were immersed in the prepared SBF solution. Following designated incubation periods (7 and 14 days), the SBF was aspirated, and the scaffolds were lyophilized. The resulting samples were then subjected to X-ray diffraction (XRD) and scanning electron microscopy (SEM) analysis to characterize the nanoHAp formation and their structural and morphological properties.

#### 2.5) Drug release profile for Bioglass embedded scaffold

##### 2.5.1) Standard curve for painkillers and antibiotics

A standard curve for painkillers, paracetamol, and diclofenac was plotted by thorough the spectrometric analysis of known concentrations of the drugs at 196 nm and 273 nm respectively.

Similarly, Standard curve for antibiotics like Amoxicillin, Doxycycline, and Augmentin was plotted at 270nm, 488nm, and 270 nm.

##### 2.5.2) Drug release profile studies of Painkiller

Solutions of painkillers (Paracetamol, Aceclofenac) were prepared at concentrations of 0. 25, 0. 5, 1. 0, and 1. 5 g/L. Scaffolds integrated with bioglass were individually immersed in each solution and left undisturbed for three days for efficient and optimum coating of drugs onto the scaffolds. Subsequently, the scaffolds were air-dried at room temperature. The dried scaffolds were then transferred to PBS solution, and their UV-visible spectrometric analysis was carried out at time intervals of 30, 60, 90, 120, and 150 minutes(**Mo et al., 2023**).

##### 2.5.3) Antibiotic release profile studies

A 20 mL solution of 8% sodium alginate was prepared. To this solution, 10 mg of bioglass was added, which was capped with either 1 mg or 5 mg of antibiotic (Amoxicillin, Augmentin, Doxycycline). The mixture was stirred magnetically until a homogeneous dispersion was achieved. Following which the mixture was added with cross linked using chilled CaCl^2^ and further subjected to lyophilization. The resulting scaffolds were then immersed in PBS solution. UV absorbance readings were subsequently taken at time intervals of 30, 60, 90, 120, and 150 minutes.

#### 2.6) Cytocompatibility assay

The colorimetric MTT test was applied to assess the cytocompatibility of the cells following interaction with the scaffold samples:

- Bioglass + Alginate .
- Bioglass +Alginate + Amoxicillin.
- Bioglass + Alginate+ Amoxicillin + Paracetamol.

The MTT assay determines the extent to which viable cells can cause the reduction of the tetrazolium salt of MTT. The assay was carried out on MG 63 (Osteosarcoma cell lines), L929 (Mouse fibroblast cell lines), and POB (Primary Osteoblast cells) were cultured under conventional culturing conditions.

Following a seeding density of 104 cells per well into 96-well plates, under aseptic circumstances, the Microspheres with nanoparticles, antibiotics, and painkillers in different permutations and combinations were left immersed in full medium for 48 hours at 37 ◦C with agitation. The medium containing the leachable was collected in a Falcon tube following the incubation period(**Zhou et al., 2020**)

#### 2.7) FTIR

The FTIR analysis was carried out for Bioglass, Bioglass embedded alginate scaffold, and Nanohydroxyapatite embedded alginate scaffold integrated with paracetamol and antibiotic. FTIR was carried out using a Perkin Elmer Frontier FTIR spectrophotometer operating in the wavelength range 4000-600 cm^-1^ to study whether the chemical integrity of the bioglass alginate scaffold was interfered with by the drugs integrated with the scaffold. The method followed was the ATR method(**Zapata et al., 2021**).

#### 2.8) ALP activity

The alkaline phosphate activity (ALP) was done to analyse the biological effects of the scaffold. Cells comprising odontoblast-like cells (OLC), primary osteoblasts (POB), and their co-culture (OLC+POB), were seeded onto pre-sterlized scaffolds and maintained under standard sustained growth media. At particular time intervals (Day 7 , 14, 21), cells were subjected to enzymatic detachment from the scaffolds using 0. 25% trypsin-EDTA solution to ensure efficient adherent cells. The collected cell suspensions were subjected to DNA isolation using the standard genomic DNA extraction procedure to obtain high-quality nucleic acids. The extracted DNA was quantified spectrohotometrically to estimate concentration and purity , which served as an indirect measure of cell proliferation and viability at each time point (**Haider et al., 2023**).

#### 2.9) Statistical Analysis

Experiments were carried out in triplicates and results were represented in the form of mean ± standard error. Data analysis and graphical representations were performed using Microsoft Excel 2019 and OriginPro 9. 0.

### 3) Results and Discussions

#### 3.1) Bioglass Synthesis and Characterization

The synthesized Bioglass had a characteristic greyish colour **(Figure 1)**. The Bioglass was subjected to XRD, which confirmed the bioglass synthesis. The bioglass is amorphous in nature. Sharp peaks at 2 theta values of 26. 6□ , 36. 5 □, 40. 3 □, 45. 8□ indicate cristobalite formation(**Kim et al., 2013**). Peak at 31. 8, 29. 9 indicates the formation of Pseudowollastonite(**Chandrasekar et al., 2013**). Cristobalite has a wide range of dental applications, while pseudowollastonite has bone regeneration properties. Peak at 32⁰ indicates the formation of Tricalcium Phosphate (Ca3(PO4)2)( JCPDS card 9–348 alpha-TCP)**(Figure 2)**. From the SEM analysis, we observed that the nanoparticles were amorphous in nature and spherical in shape. The nanoparticles were in the range of 154. 2nm to 227. 9nm in size **(Figure 3)**. The agglomerations found in the bioglass nanoparticles can be due to Ostwald ripening (**Ravarian et al., 2010**).

**Figure 1:**
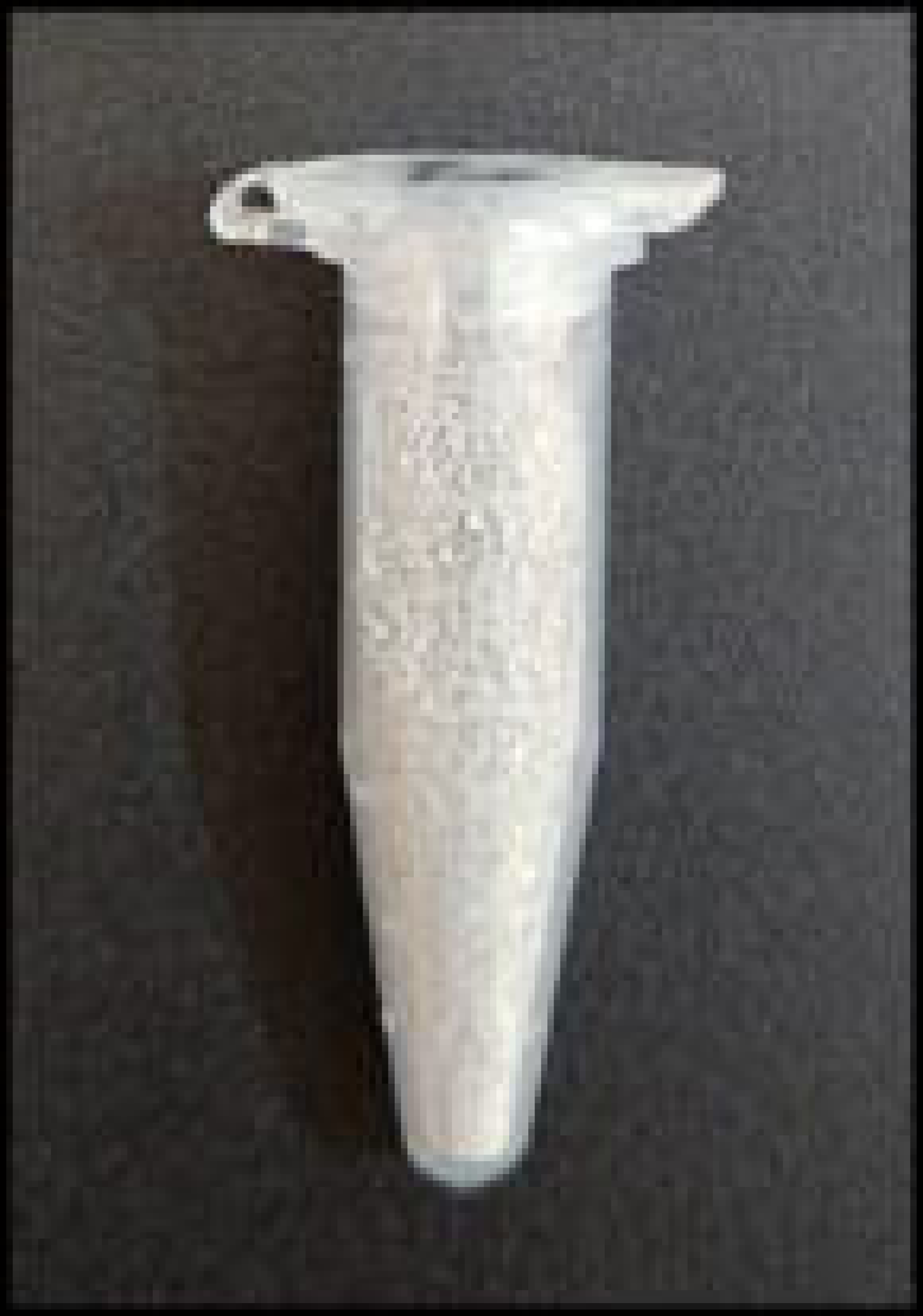
Bioglass

**Figure 2:**
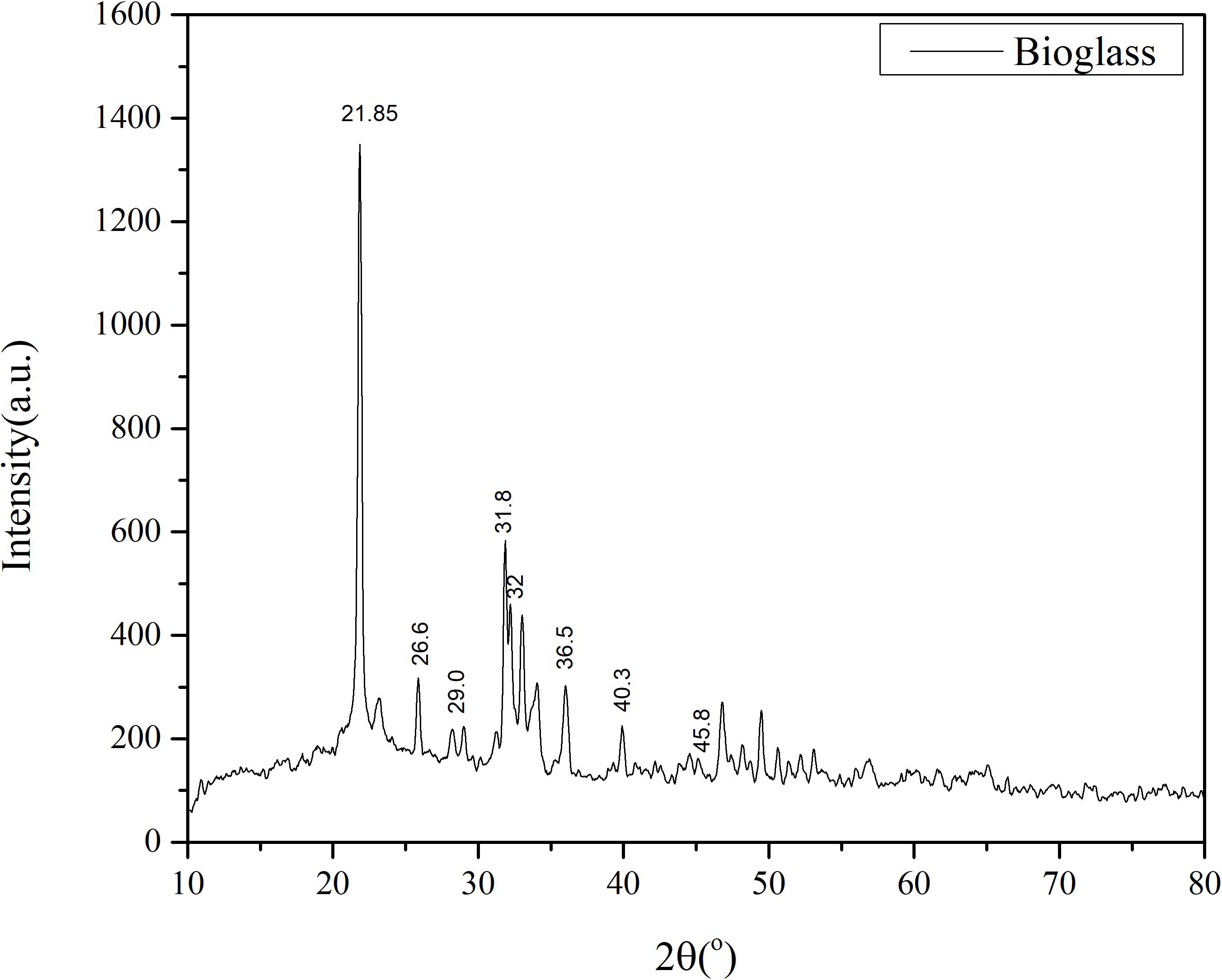
XRD result of Bioglass.

**Fig. 3:**
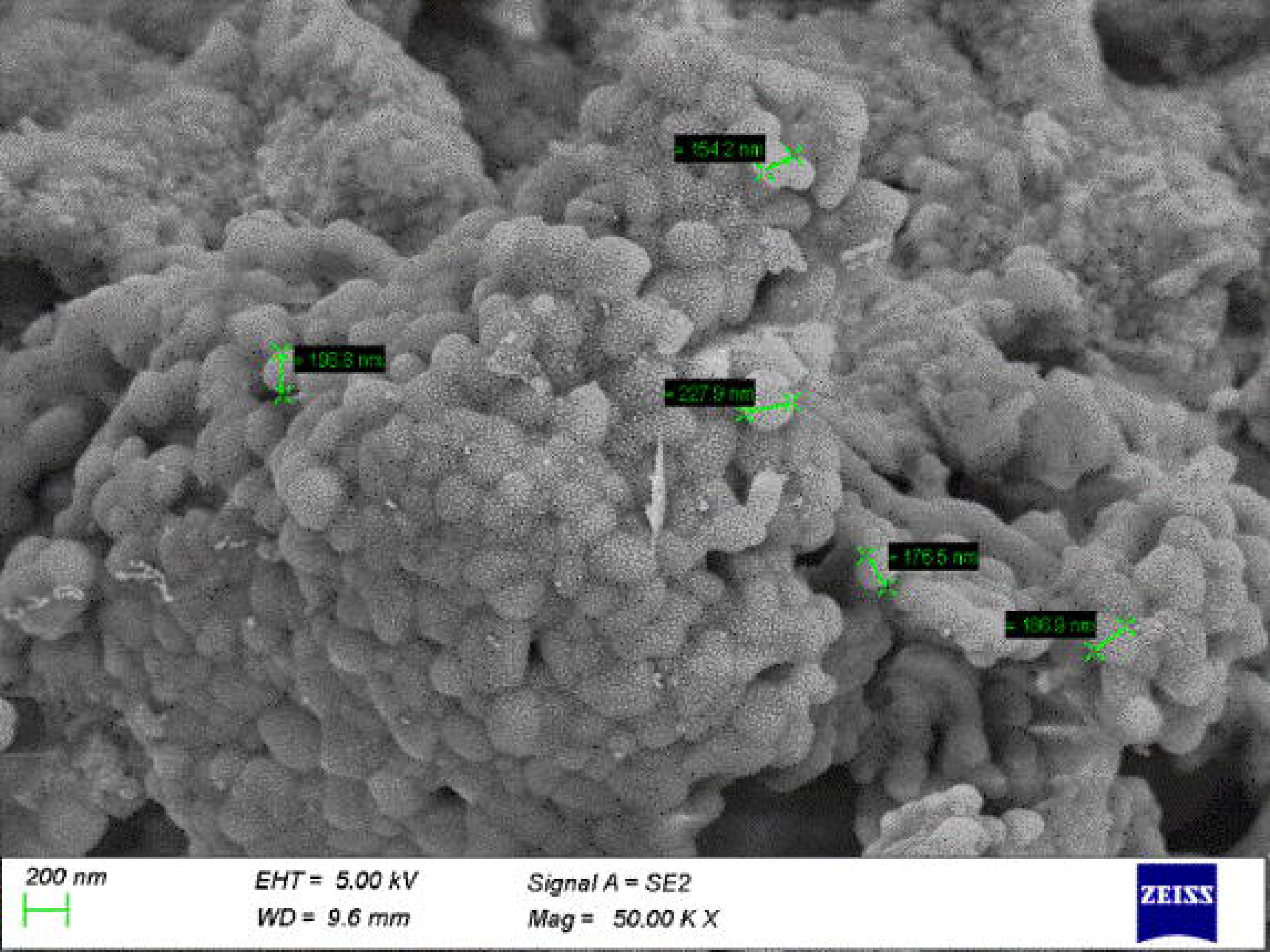
SEM analysis of bioglass

#### 3.2.) Preparation and characterization of the Bioglass embedded scaffold

The prepared bioglass-embedded alginate was found to be sturdy (**Figure 4**). From the XRD results, it was confirmed that the bioglass was embedded in the alginate without undergoing any changes. From the XRD, we can infer that the bioglass has been integrated into the alginate without losing its chemical integrity, as peaks of Bioglass were found to be retained in the bioglass-embedded alginate scaffold **(Figure 5)**. From the SEM analysis, it was found that the bioglass was successfully embedded onto the surface of the alginate **(Figure 6)**.

**Figure 4:**
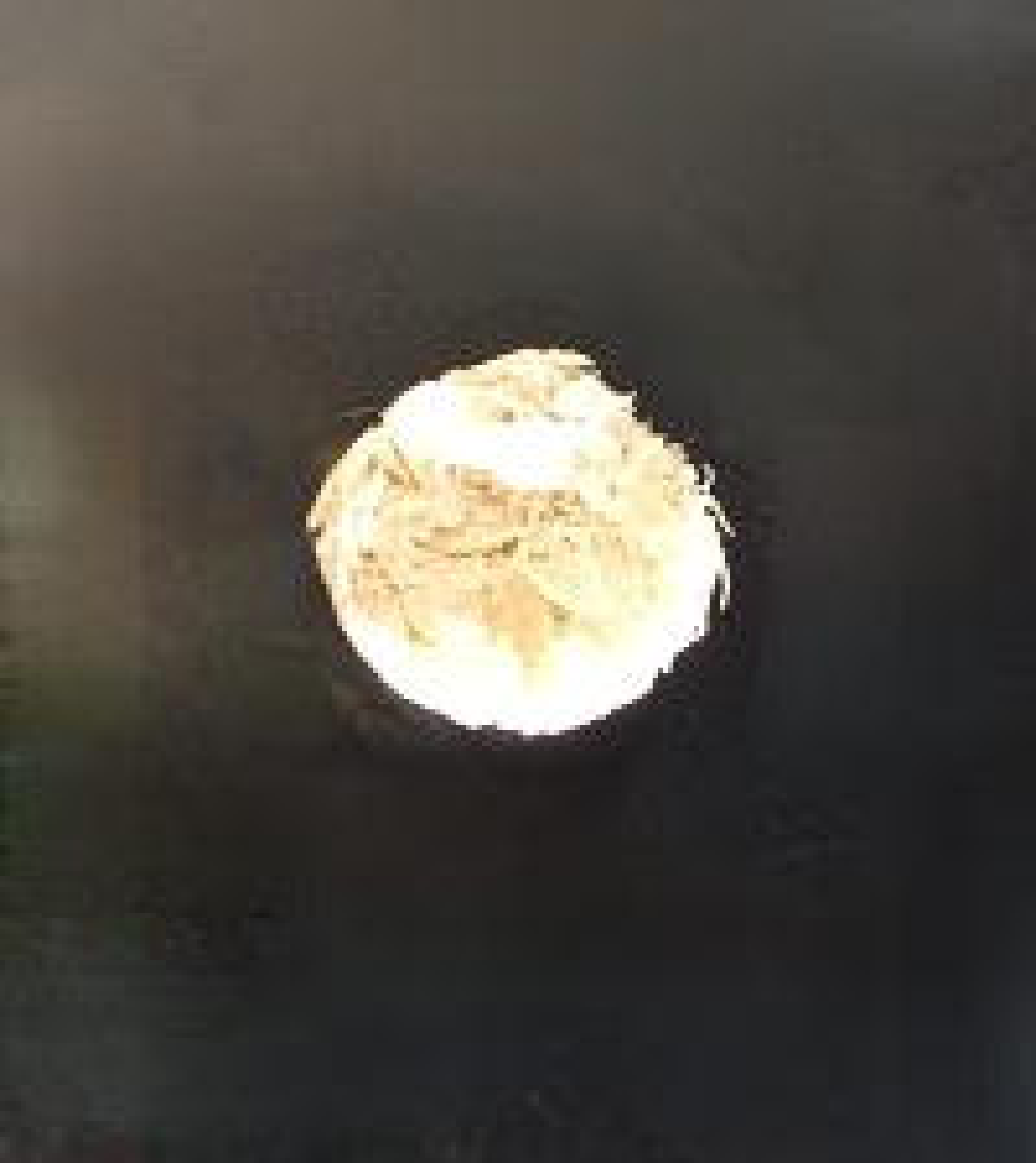
Bioglass integrated scaffold.

**Figure 5:**
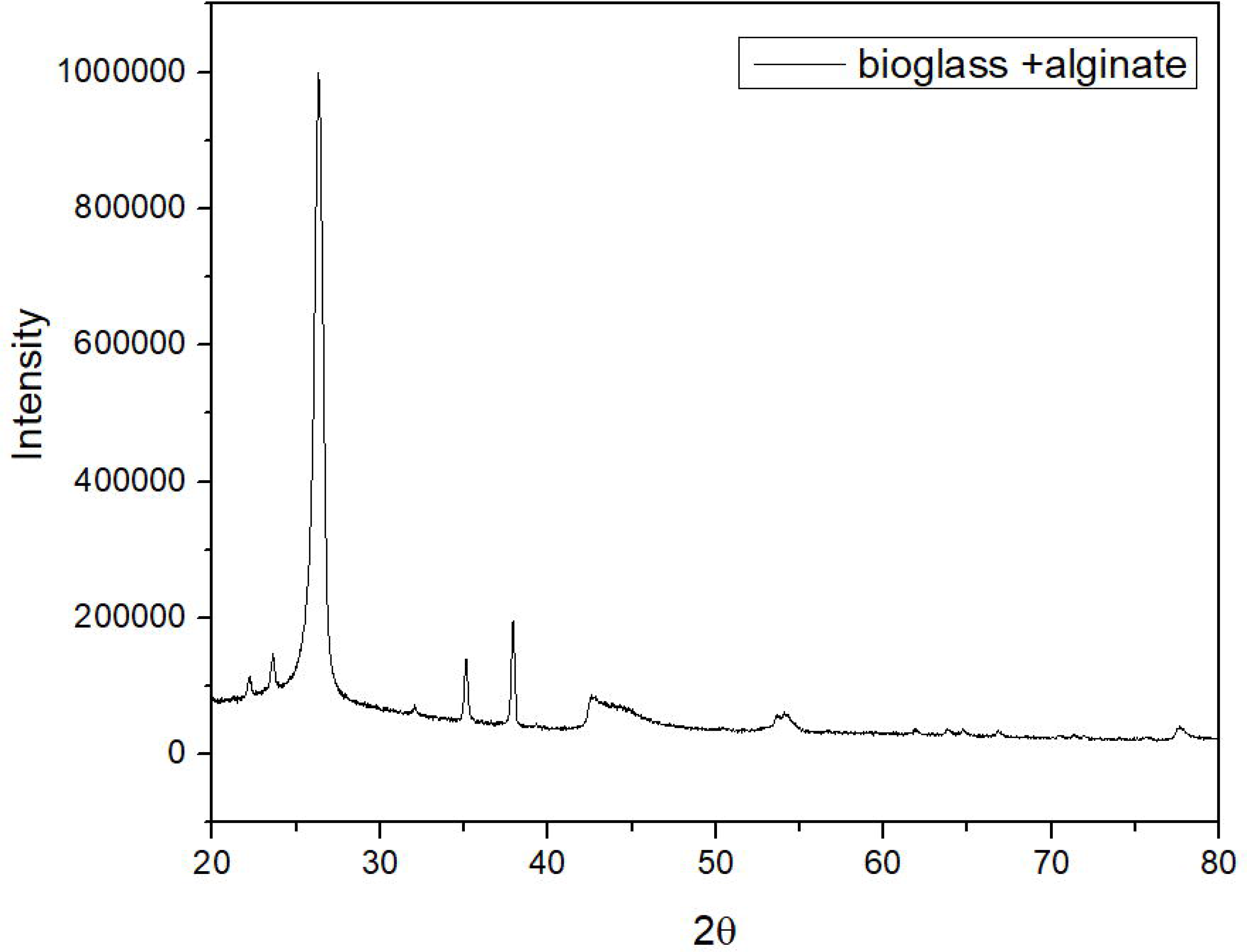
XRD results of bioglass integrated scaffold.

**Figure 6:**
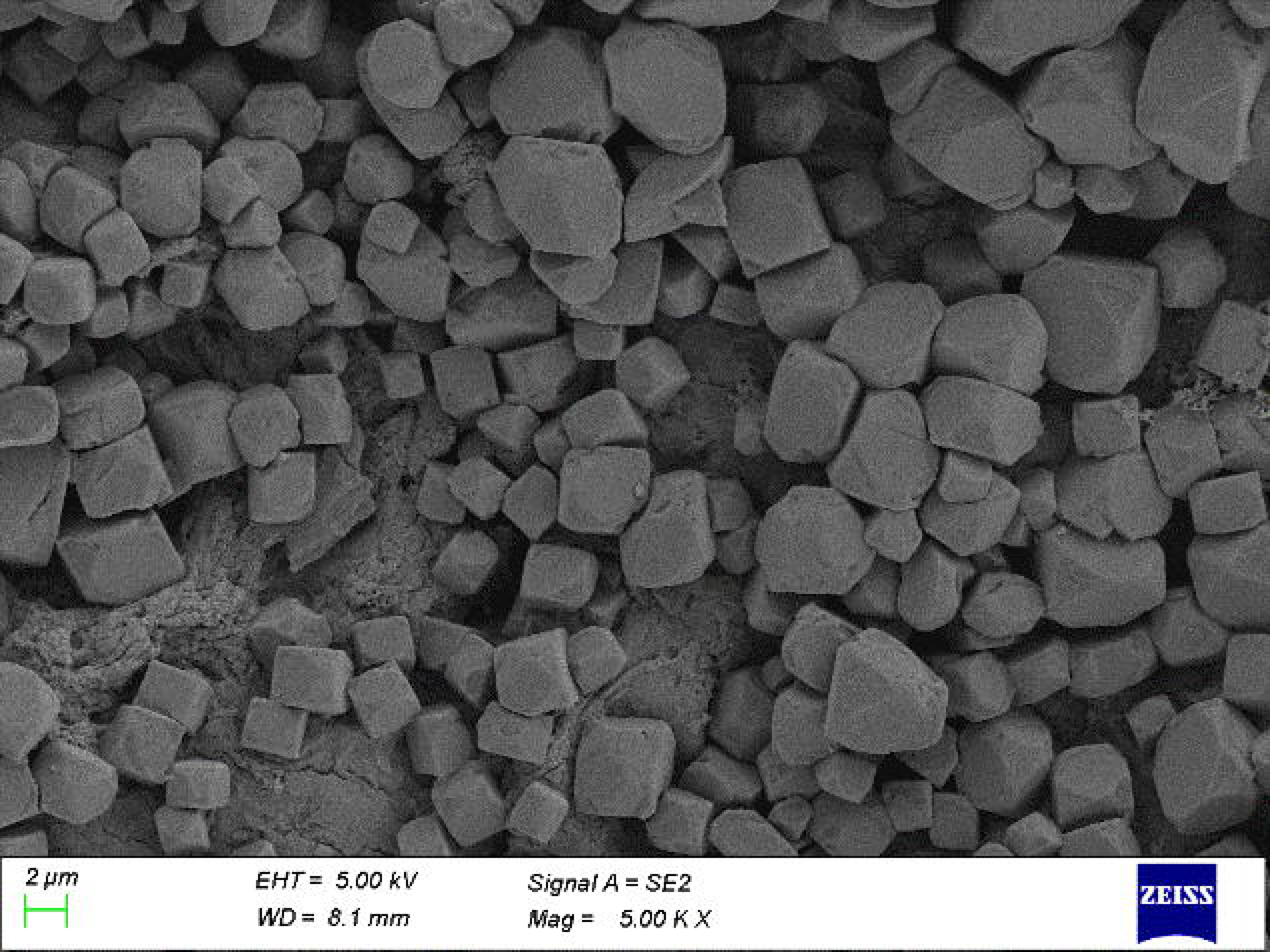
SEM analysis results of bioglass-embedded alginate scaffold.

#### 3.3) Swelling and degradation analysis of bioglass-embedded scaffolds

##### 3.3.1) Swelling analysis

Swelling analysis was carried out to understand the fluid uptake ability and the mechanical properties of the scaffold when alginate was integrated with the bioglass in comparison to just the alginate to validate the influence of the integration of bioglass on impacting the scaffold’s performance. The Swelling studies were carried out as explained earlier. For both the scaffolds alginate (control) and the bioglass embedded alginate scaffold (test) , the maximum swelling percentage was observed on Day 11, after which it was observed that there was a significant decrease in wet weight of the control. In contrast the wet weight decrease was gradual in the test till day 21 . The maximum swelling was observed on Day 11 for the bioglass embedded alginate scaffold, it was found that the maximum swelling percentage was observed in the scaffold embedded with bioglass (1628. 28±0. 09%) in comparison to just alginate (1152. 94±0. 05 % ). Hence, we can infer that the bioglass-embedded alginate scaffold has better swelling efficiency. In comparison to the control scaffold **(Figure 7)**

**Figure 7:**
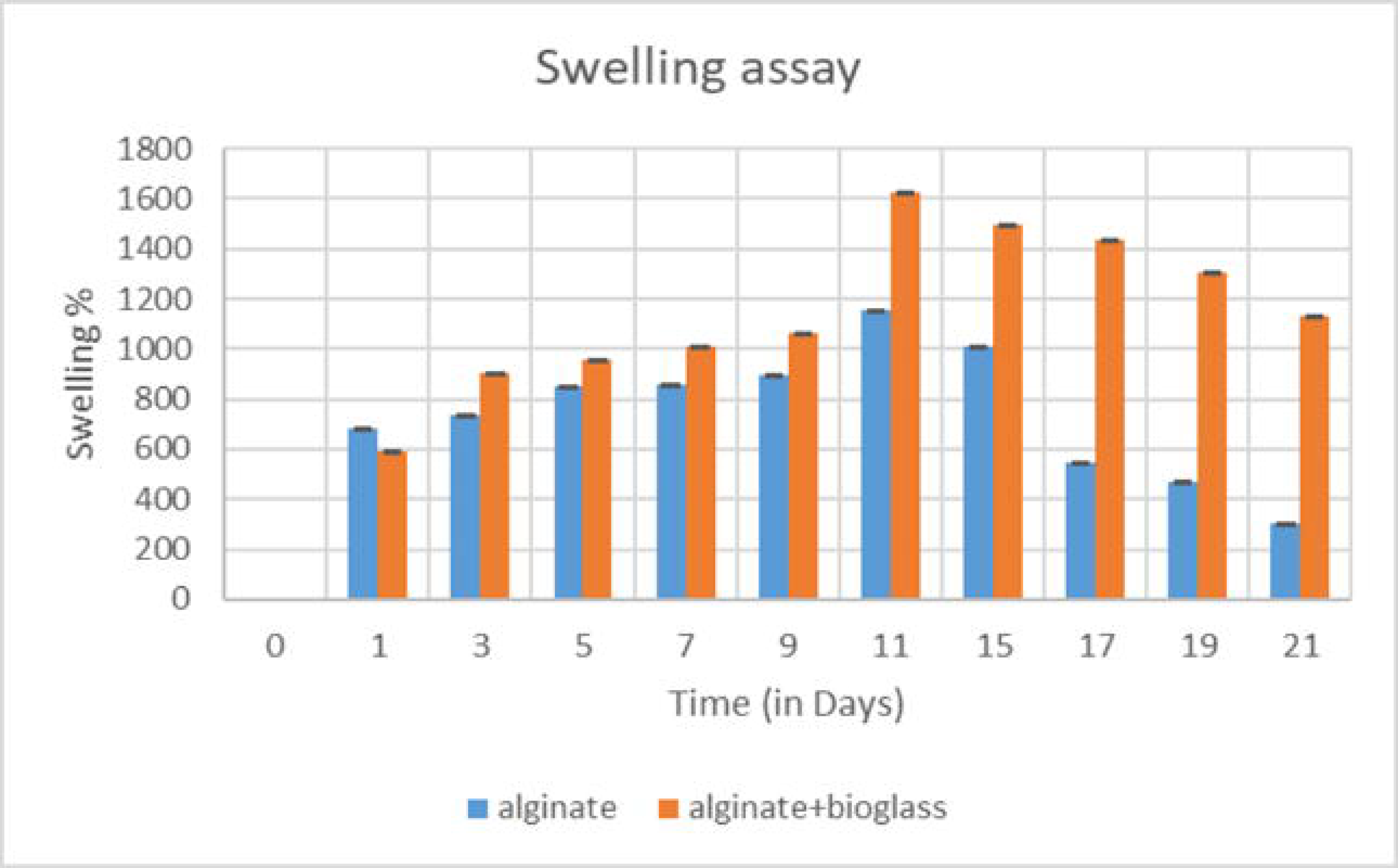
Swelling assay Control(alginate), Test(alginate+bioglass)

##### 3.3.2) Degradation assay for bioglass-embedded scaffold

Scaffold design is a vritical parameter in tissue engineering design. Optimally, the degradation kinetics of the scaffold should be closely matched to the rate of neotissue formation, thereby facilitating appropriate structural integration and functional maturation of the regenerated tissue. From the degradation assay, it was inferred that the test scaffold degradation was slow in comparison to the test. While the wet weight increase was observed on day 1 the wet weight it was observed from day 3 that the wet weight of both scaffolds was observed to decrease. While the decrease of wet weight in the test scaffold was gradual and consistent, it was observed that for the control, there was a sharp decrease in wet weight from day 9 to Day 15 **(Figure 8)**.

**Figure 8:**
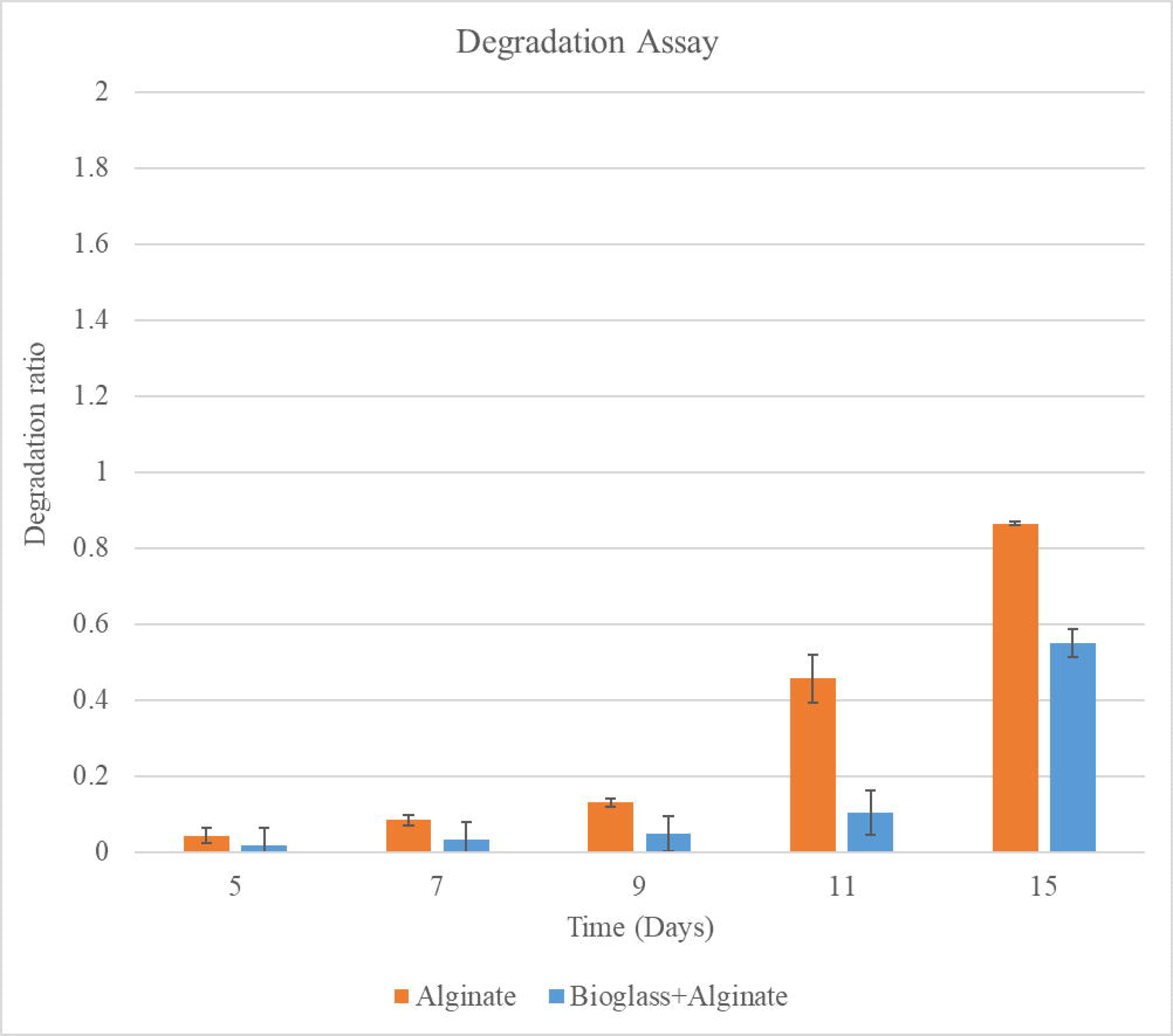
Degradation assay of BG-embedded scaffold.

#### 3.4) Biomineralization studies

Biomineralization studies were conducted to assess the scaffold’s ability to enhance the formation of bone-like apetite on its surface, thereby confirming its suitability for bone regeneration applications. The hydroxyapatite formation was observed as the peaks were evident at 2θ values of 25. 9□, 31. 8□, 32. 2□, 34. 0□, 39. 8□, 46. 7□, 49. 5□ (JCPDS no. 09-0432). It was observed that the peaks of bioglass shifted as the intensity of the bioglass peaks decreased. This was due to amorphous bioglass reacting and being converted to HAp. From the SEM images of the bioglass embedded alginate scaffold, which was immersed in SBF for 14 days, we observe an appreciable level of hydroxyapatite deposition on the scaffold, indicating that the bioglass integration into the scaffold enhances the hydroxyapatite deposition and is suitable for bone tissue engineering applications (**Figure . 9**).

**Figure 9:**
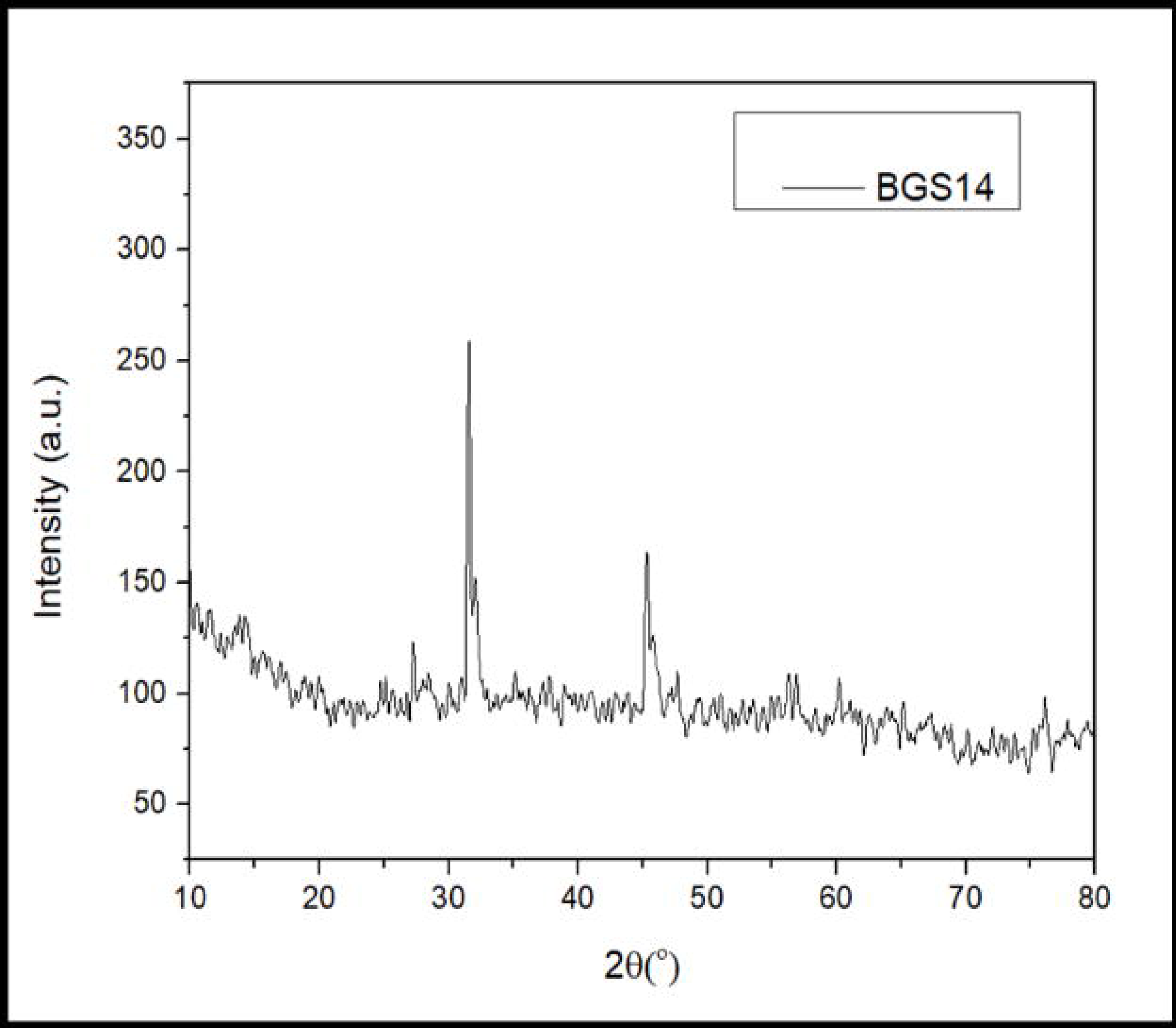
XRD results for scaffold embedded with Bioglass, immersed for 14 days in SBF.

**Figure 10:**
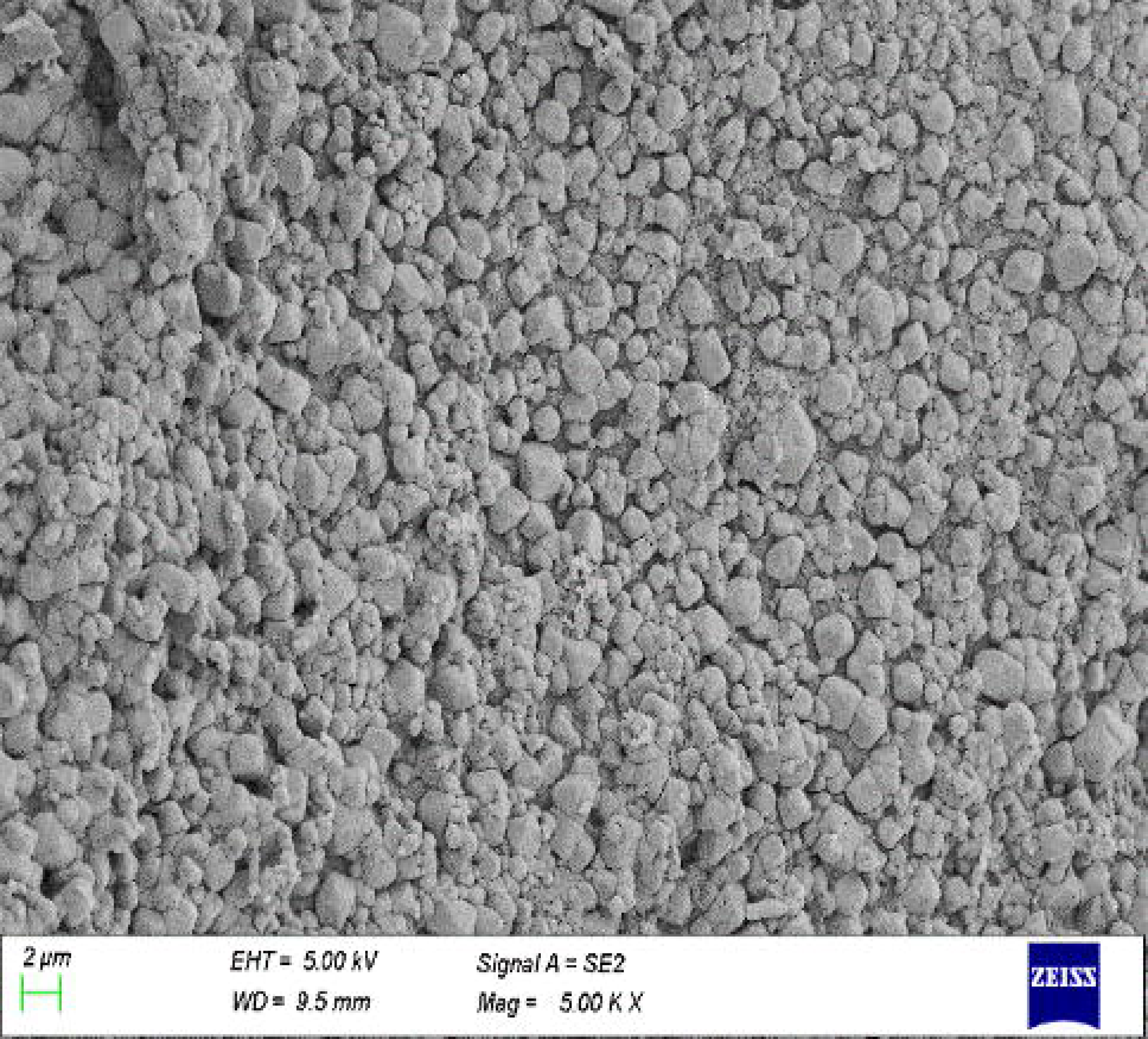
SEM image for scaffold embedded with Bioglass, immersed for 14 days in SBF.

**Figure 11:**
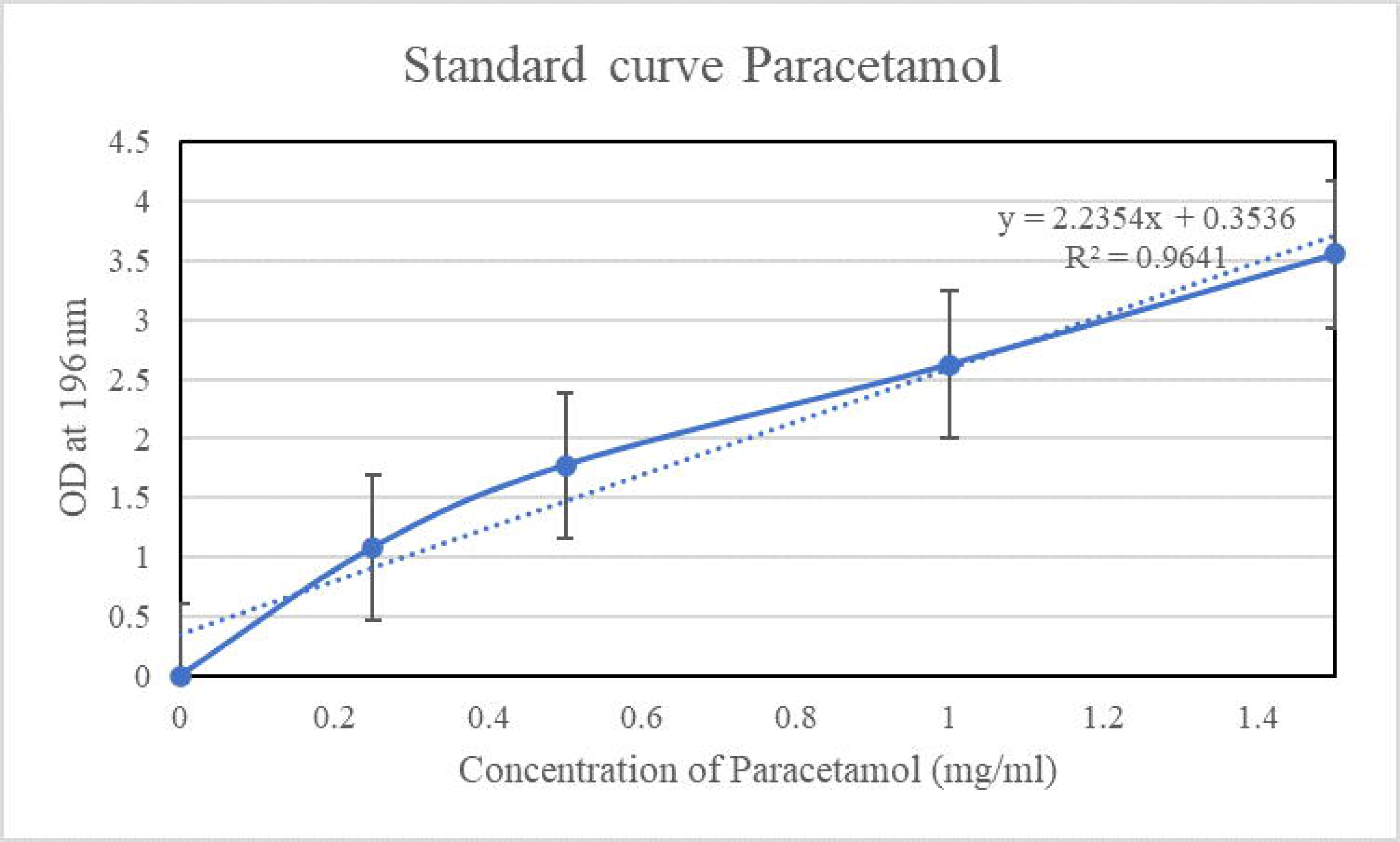
Standard curve for Paracetamol.

#### 3.5) Drug release profile studies

##### 3.5.1) Standard curve for painkillers and Antibiotics

The standard curves were prepared according to the procedure stated in the methodology for both the painkillers (Paracetamol and Aceclofenac) and the Antibiotics (Amoxicillin, Doxycycline, and Augmentin)(**Figure 12-15**).

**Figure 12:**
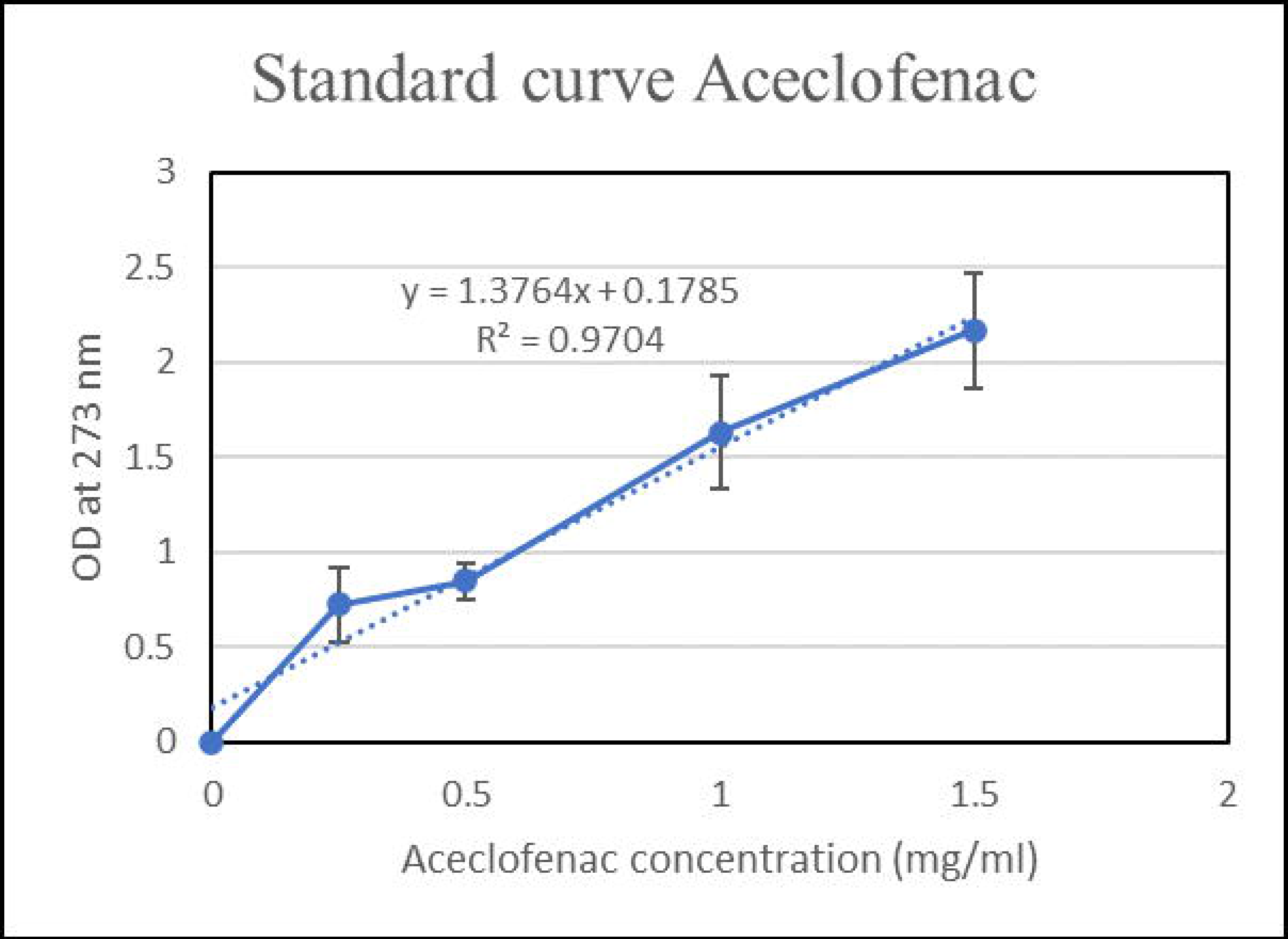
Standard curve Aceclofenac.

**Figure 13:**
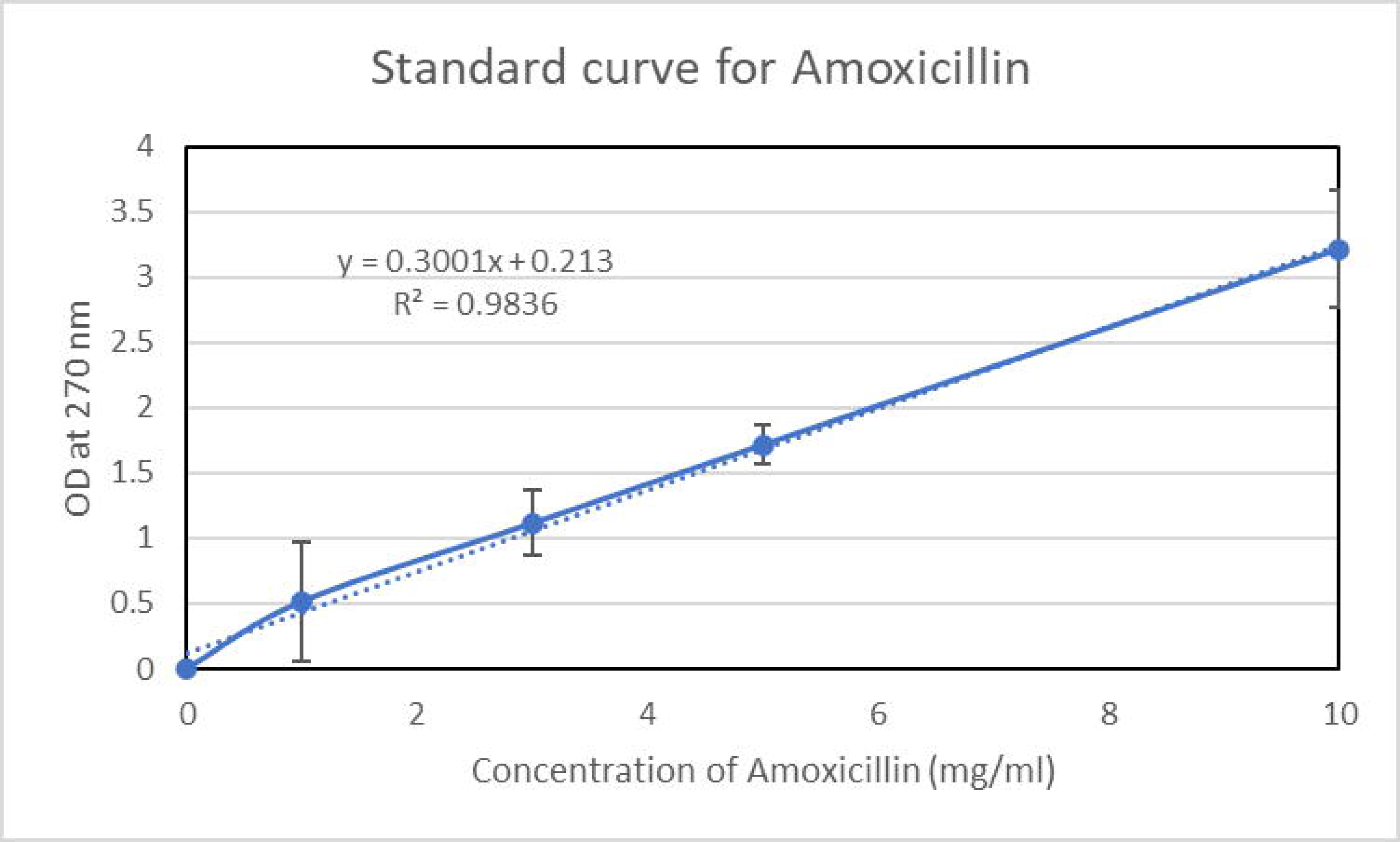
Standard curve of Amoxycillin.

**Figure 14:**
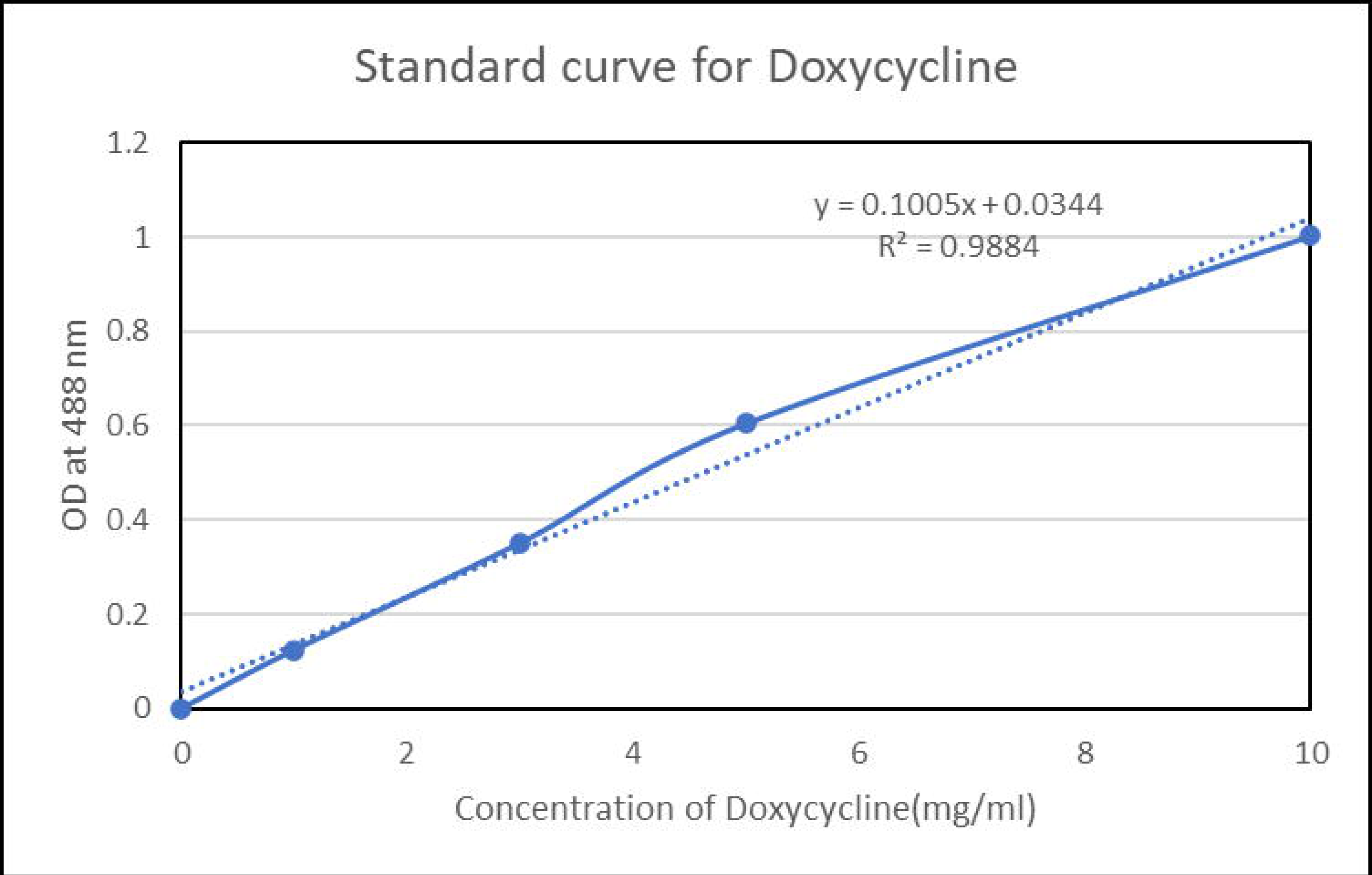
Standard curve of Doxycycline.

**Figure 15:**
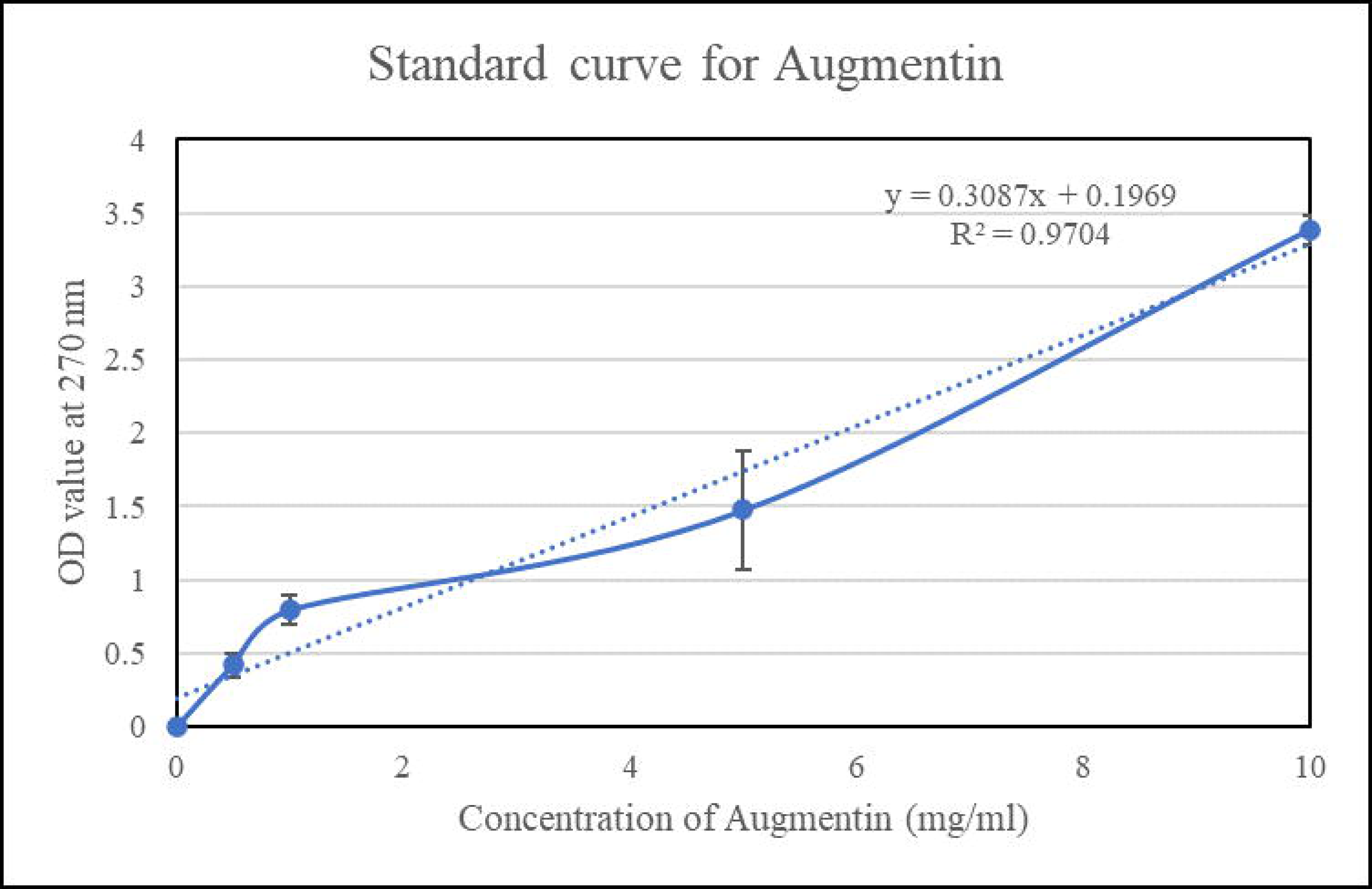
Standard curve of Augmentin.

##### 3.5.2) Painkillers release profile

From the Painkiller release profile studies, it was observed that in a scaffolds with Aceclofenac as well as Paracetamol, a rapid release was observed. The maximum release of the painkillers was at the 90^th^ minute, and in both cases for the scaffold with 1. 5 mg/ml of paracetamol/ aceclofenac concentration. When we compared the scaffold with Paracetamol and with Aceclofenac, it was noted that the scaffold with Paracetamol had a better release in comparison to the scaffold with Aceclofenac as the scaffold integrated with 1. 5mg/ml of paracetamol gave a maximum release of 1. 452±0. 007mg/ml of paracetamol while the scaffold integrated with 1. 5 mg/ml of Aceclofenac gave a maximum release of 1. 246 ±0. 002 mg/ml at 90^th^ minute . Overall, the results of the painkiller release profile studies are in synchronisation with our concept of wanting a burst release of painkiller from the scaffold to treat the dental bone injury**(Figure 16)**.

**Figure 16:**
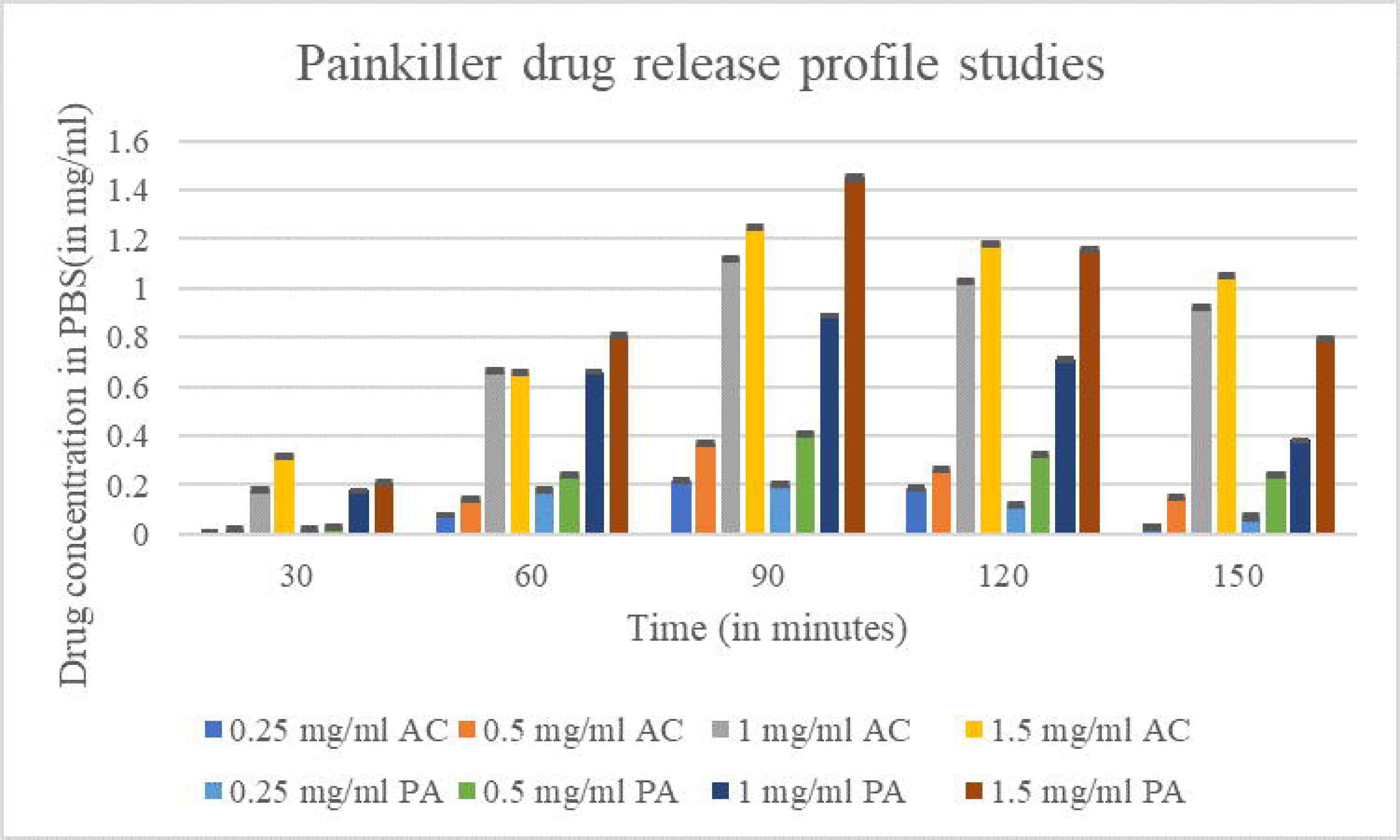
Painkiller drug release profile.

##### 3.5.3) Antibiotic release profile studies

From the Antibiotic release profile studies, it was observed that there was sustained release of Antibiotics from the scaffold, which was due to capping of the antibiotic onto the bioglass. This sustained release is beneficial for our dual drug release scaffold. it was observed that for the scaffolds integrated with 1 or 5 mg/ml of Amoxicillin and Doxycycline the maximum release was observed to be at 1500^th^ minutes However in the case of scaffold integrated with Augmentin , the scaffold integrated with 5 mg/ml of Augmentin showed maximum release at 1440^th^ minute while the scaffold integrated with 1mg/ml of Augmentin showed maximum release at 1500^th^ minute. Considering the overall antibiotic release profile scaffold integrated with 5 mg/ml of Amoxicillin was found to give maximum release (3. 95±0. 004 mg/ml) at 1500 minutes in comparison to the scaffolds integrated with Doxycycline and Augmentin **(Figure 17)**.

**Figure 17:**
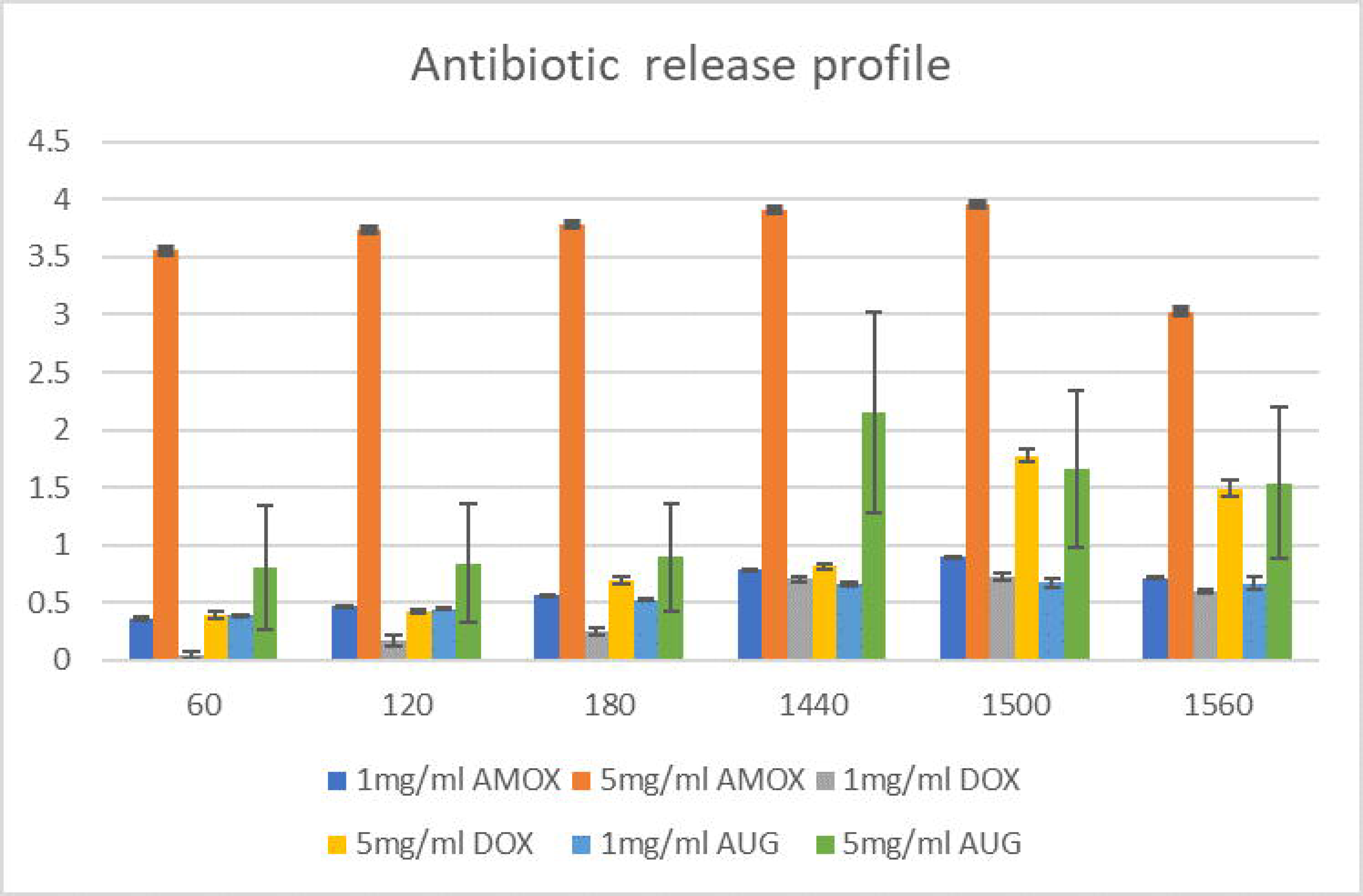
Antibiotic release profile studies.

#### 3.6) Cytocompatibility studies

Cytocompatibility studies were carried out on osteoblastic cell lines such as MG63 and POB, along with L929, which is a fibroblastic cell line, to understand the compatibility and efficacy of the prepared scaffold for dental bone tissue engineering applications. It was found that on average, the cell lines were 98% compatible with the scaffolds prepared (bioglass alginate scaffold, bioglass alginate scaffold integrated with Amoxicillin, Bioglass embedded alginate scaffold integrated with Amoxicillin and Paracetamol), making the scaffold highly suitable for dental bone tissue engineering applications(**Figure 18**).

**Figure 18:**
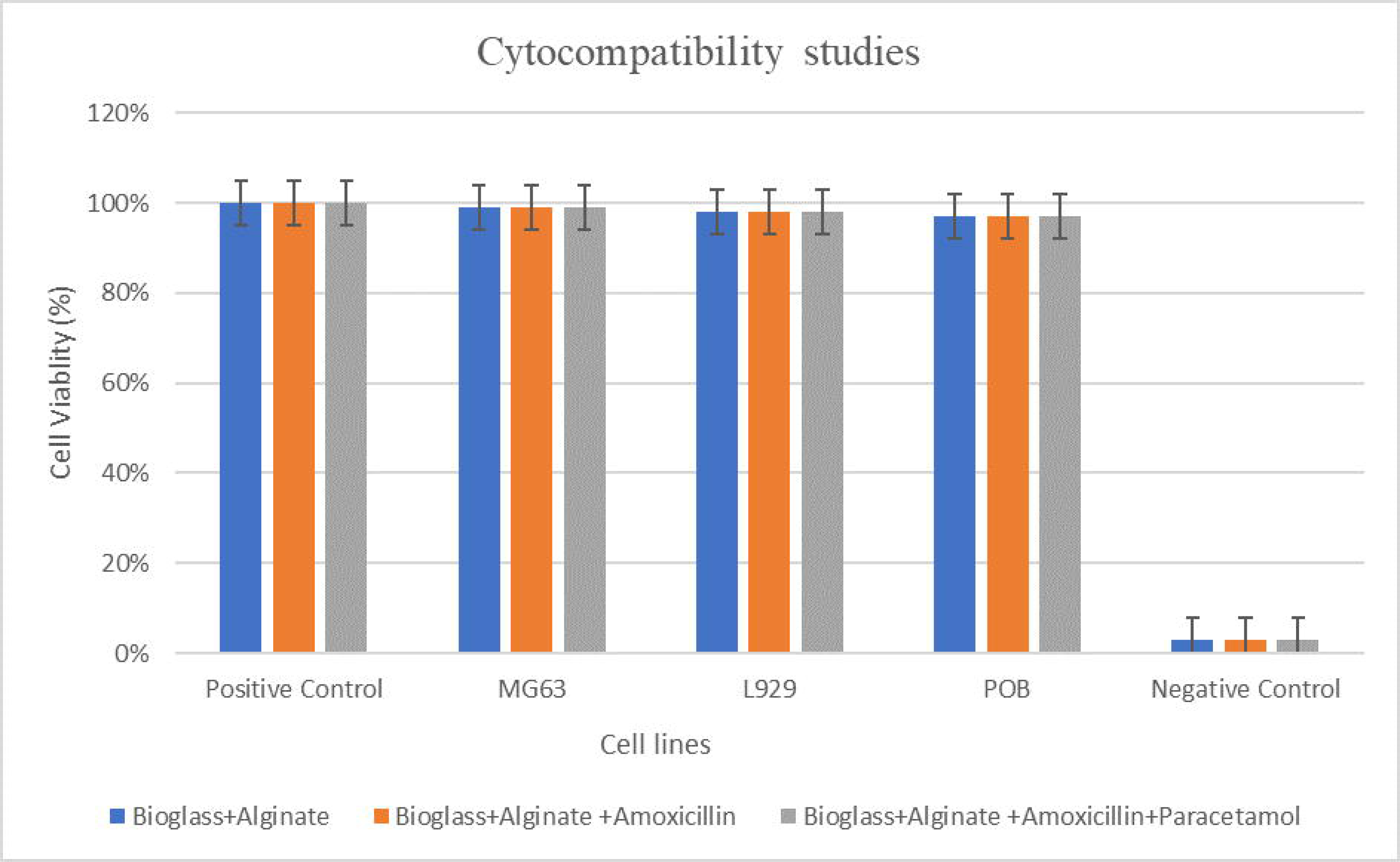
Cytocompatiblity studies.

#### 3.7) FTIR analysis

FTIR spectroscopy was carried out to confirm that the integration of the drugs did not interfere with the chemical integrity of the bioglass-embedded alginate scaffold. Broad, intense peaks at the region 3766–3369 cm□^1^ were correlated to overlapping OH and NH stretching of alginate hydroxyls, bioglass silanols, and drug amide groups, showing an extensive hydrogen-bonding network throughout the matrices. Specific carboxylate bands (COO□ asymmetric/symmetric stretching at 1598–1624 and 1430–1423 cm□^1^), along with CO stretching/OH bending near 1300–1317 cm□^1^ and COC/CO vibrations at 1148, 1116, 1080 and 1027 cm□^1^, validated the retention of the alginate backbone and its ionized carboxylate functionalities after integration of bioglass and drugs, while the NH bending/aromatic band at 1500 cm□^1^ confirmed the presence of intact amoxicillin or paracetamol. In the lower wavenumber region, characteristic Si–O–Si stretching and bending bands at 1080, 944–938, 882–809, and 672–658 cm□^1^ confirmed an amorphous silicate–phosphate bioglass network comprising non-bridging oxygens and surface Si–OH groups having the potential of interacting with alginate and drug molecules. The partial overlap of C–O and Si–O–Si bands approximately at 1116–1080 cm□^1^, along with composite-specific overtone or combination bands between 2322 and 1980 cm□^1^, showing strong hydrogen-bonding and electrostatic interactions between alginate carboxylates, bioglass silanols, and drug functional groups, without the appearance of new covalent-bond peaks that would indicate drug degradation. Overall, these features substantiate that the scaffolds consist of structurally intact bioglass–alginate networks in which amoxicillin and paracetamol are physically entrapped and non-covalently bound, providing a robust molecular basis for sustained, localized therapeutic release in bone-tissue engineering applications(**Figure 19, 20**)(**Sartori et al., 1997, Badita, et al 2020, Costa et al ., 2021, Bakr et al 2025, Mallah et al., 2015**).

**Figure 19:**
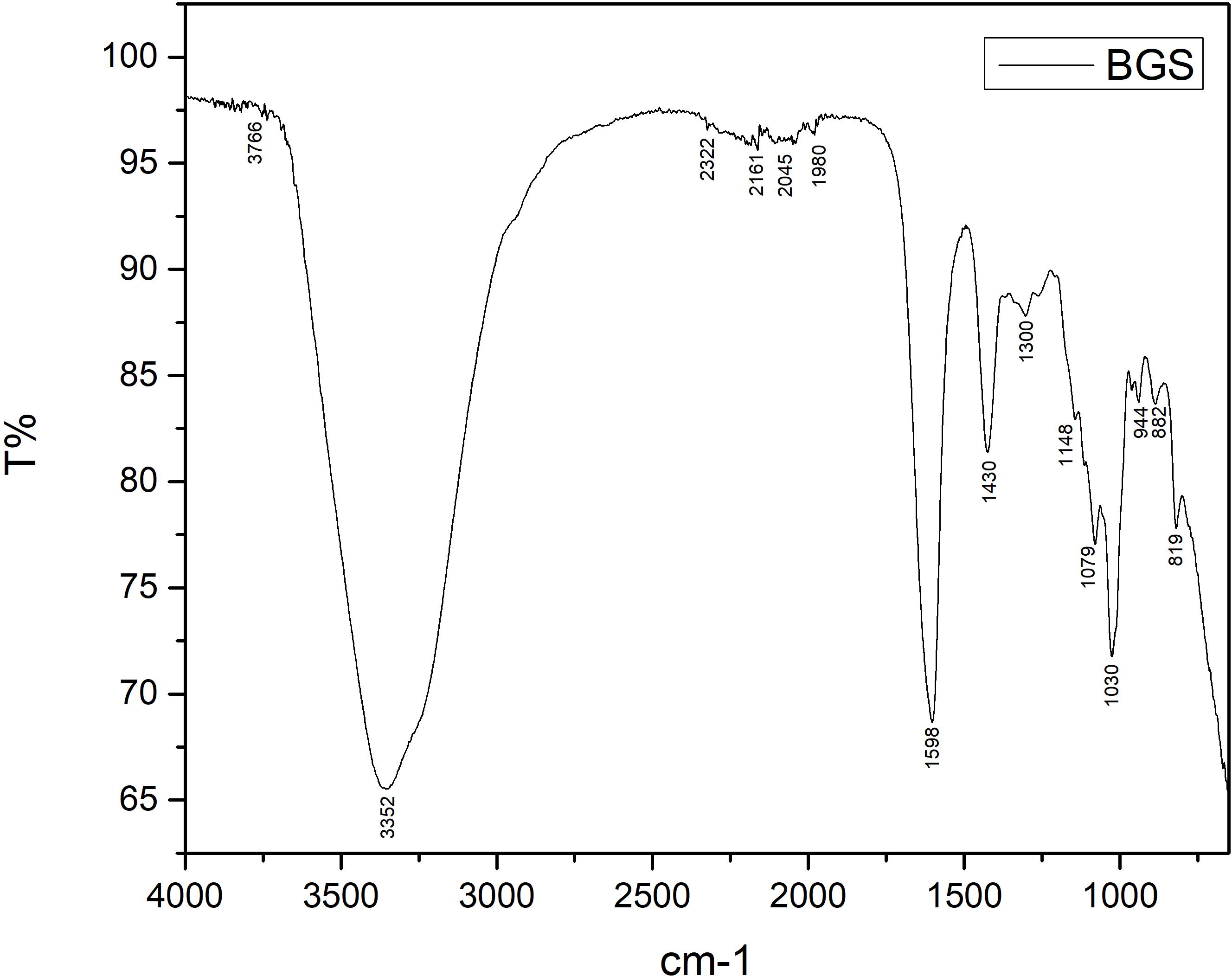
FTIR results of Bioglass scaffold.

**Figure 20:**
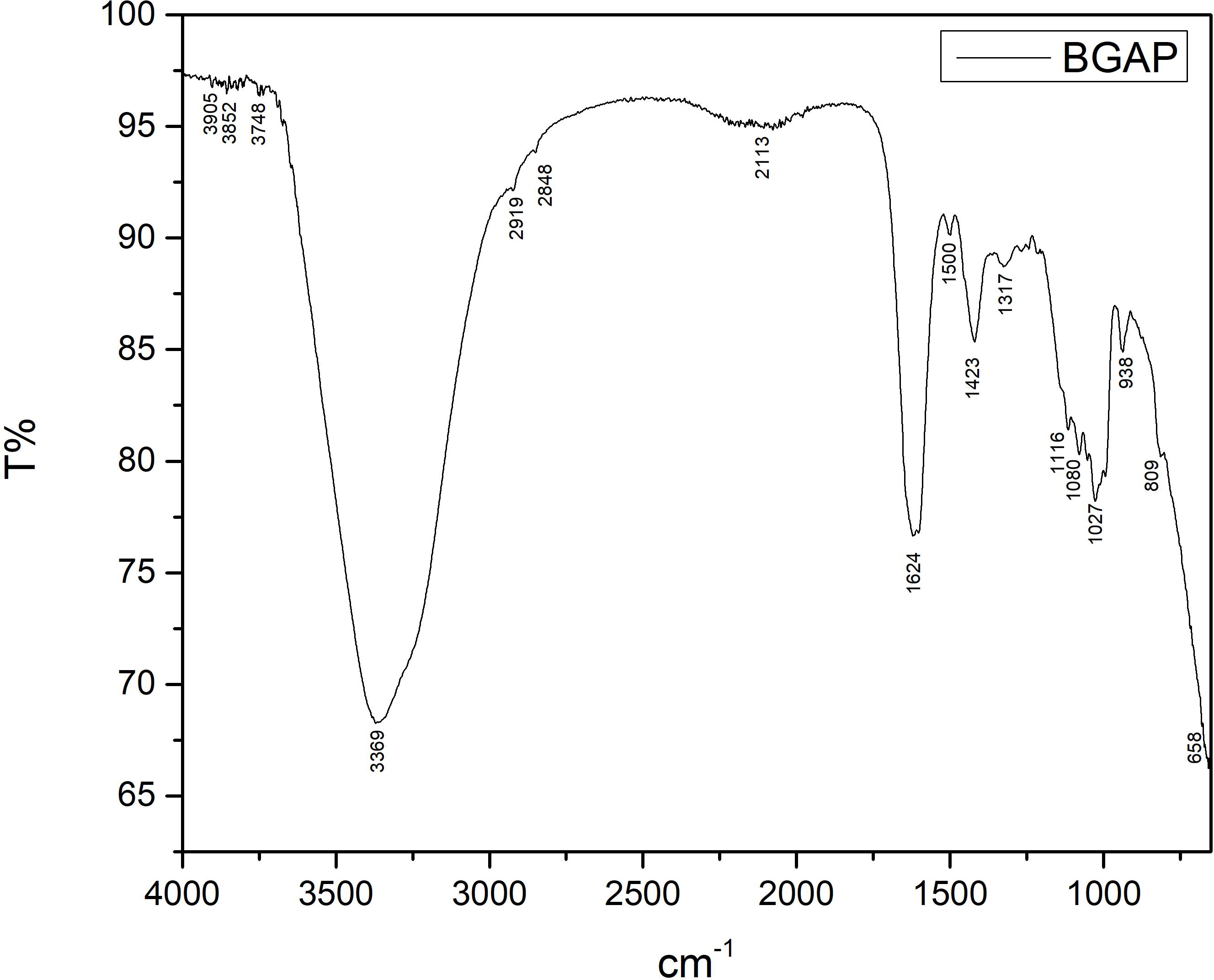
FTIR results of Bioglass embedded alginate scaffold with Amoxicillin and Paracetamol.

**Figure 21:**
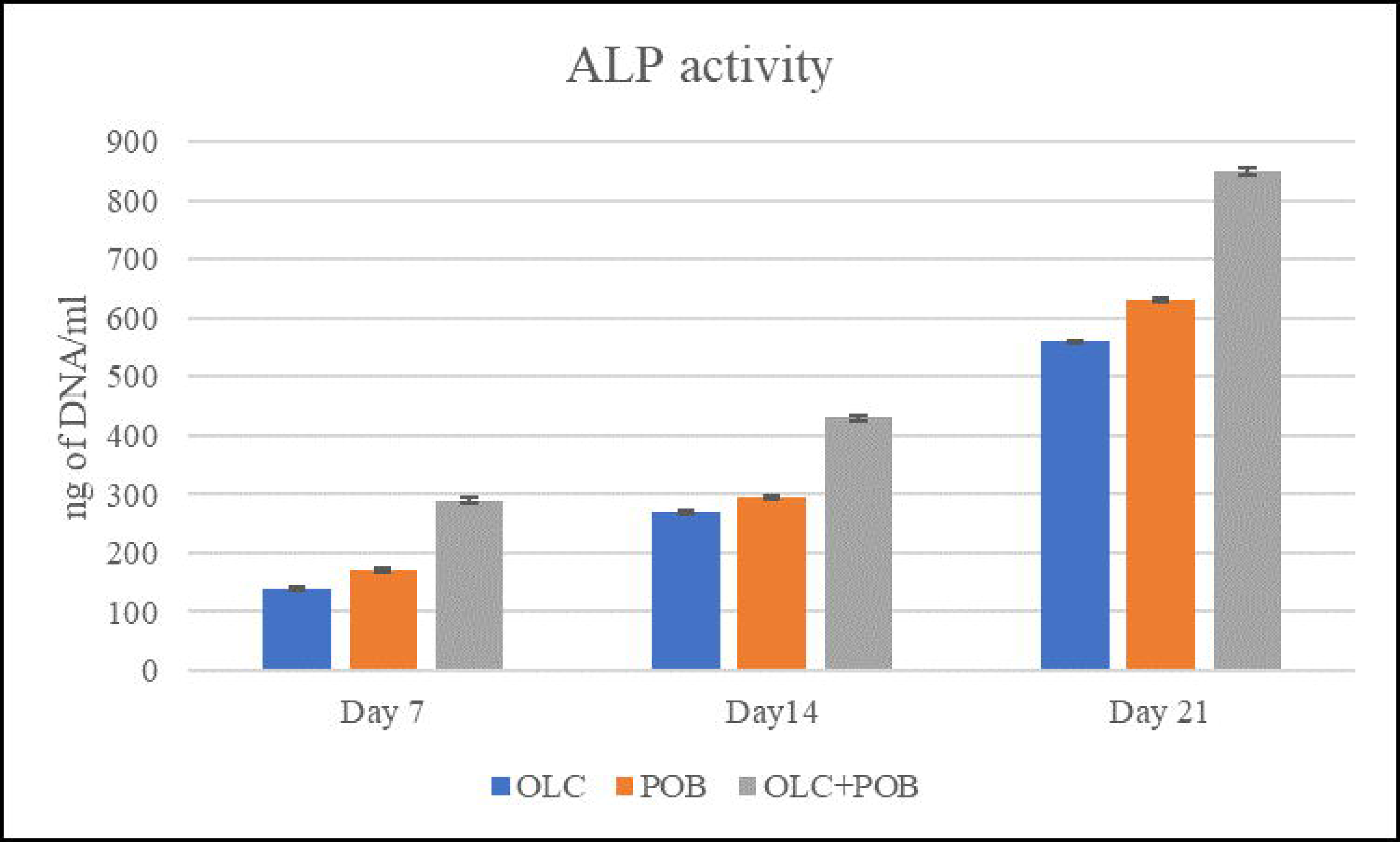
Results of ALP assay.

#### 3.9) ALP assay

The ALP activity graph Shows a time-dependent enhancement in osteogenic differentiation across all tested groups, with the co-cultured OLC+POB group uniformly observed to have higher ALP activity relative to the single cell lines (OLC and POB) at all time points. On day 7, ALP levels are modest for all groups, but by day 14 and day 21, an evident increase is observed, specifically for the co-culture, which achieves the greatest activity by day 21. This trend shows that the scaffold leachate treatment successfully promotes osteoblastic activity and matrix maturation, with synergistic effects apparent in the mixed cell population. These inferences indicate that co-culturing OLC and POB enhances osteogenic response—likely due to cell-cell interactions—and demonstrates the capability of the scaffold as a potent inducer of bone tissue formation. The results are consistent with the expected biological pattern of ALP activity as an early marker for osteoblast differentiation and indicate the added value of using co-culture systems for assessing novel biomaterials in regenerative applications

### 4) Conclusion

Traumatic dental injuries makup a major global health concern across different age groups, highlighting the need for personalized therapeutic approaches. Dental tissue enginnering provides patient specific solutions to tackle complex clinical challenges. This research reports the development of a nanocomposite scaffold with dual drug delivery functionality for dental bone tissue engineering. The scaffold was engineered to enable rapid painkiller release for immediate inflammation control and sustained antibiotic delivery to tackle infection. Bioglass was integrated to promote osteogenic activity and enhance bone regeneration, and its successful synthesis using the novel modified citric acid method was confirmed by X-ray diffraction analysis .

Swelling and degradation studies showed that the the integration of bioglass into the scaffold enhanced the swelling and reduced the degradation of the scaffold , supporting prolonged bone healing . Drug release studies demonstrated that controlled antibiotic release from bioglass embedded scaffold , with Amoxicillin-loaded scaffolds showing superior relese efficiency in comparison to the Augmentin and Doxycycline loaded scaffolds, while Paracetamol-coated scaffolds exhibited rapid analgesic release and much more efficient than the scaffolds coated with Aceclofenac. Cytocompatiblity evaluated by MTT assay showed approximately 97% viability in osteoblast and fibroblast cell line. FTIR analysis confirmed preservation of scaffold chemical integrity even on integration of drugs . Overall the scaffold shows strong potential for customizable dental bone regeneration with future in vivo validation.

